# Polarised subcellular activation of ROPs by specific ROPGEFs drives pollen germination in *Arabidopsis thaliana*

**DOI:** 10.1101/2024.01.11.575165

**Authors:** Alida Melissa Bouatta, Andrea Lepper, Philipp Denninger

## Abstract

During plant fertilisation, excess male gametes compete for a limited number of female gametes. The dormant male gametophyte, encapsulated in the pollen grain, consists of two sperm cells enclosed in a vegetative cell. After reaching the stigma of a compatible flower, quick and efficient germination of the vegetative cell to a tip-growing pollen tube is crucial to ensure fertilisation success. RHO OF PLANTS (ROP) signalling and their activating ROP GUANINE NUCLEOTIDE EXCHANGE FACTORS (ROPGEFs) are essential for initiating polar growth processes in multiple cell types. However, which ROPGEFs activate pollen germination is unknown. We investigated the role of ROPGEFs in initiating pollen germination and the required cell polarity establishment. Of the five pollen-expressed ROPGEFs, we found that GEF8, GEF9, and GEF12 are required for pollen germination and male fertilisation success, as *gef8;gef9;gef12* triple mutants showed almost complete loss of pollen germination *in vitro* and had a reduced allele transmission rate. Live cell imaging and spatiotemporal analysis of subcellular protein distribution showed that GEF8 and GEF9, but not GEF12, displayed transient polar protein accumulations at the future site of pollen germination minutes before pollen germination, demonstrating specific roles for GEF8 and GEF9 during the initiation of pollen germination. Furthermore, this novel GEF accumulation appears in a biphasic temporal manner and can shift its location. We showed that the C-terminal domain of GEF8 and GEF9 confers this protein accumulation and demonstrated that GEFs locally activate ROPs and alter Ca^2+^ signalling, which is required for pollen tube germination. We demonstrated that GEFs do not act redundantly during pollen germination and described for the first time a polar domain with spatiotemporal flexibility, which is crucial for the *de novo* establishment of a polar growth domain within a cell and, thus, for pollen function and fertilisation success.

## Introduction

Sexual reproduction is a fundamental and complex process in which male and female gametes fuse to form a zygote, which develops into an embryo. In Angiosperms, sperm cells have lost their motility, and the male gametes must be delivered to the female gametes. This is achieved by a tip-growing pollen tube formed by the vegetative pollen cell, which encloses two sperm cells. This pollen tube grows from the papilla cells of the stigma on the flower surface into the transmitting tract toward the female gametophyte inside the ovary. After reaching the female gametophyte, the pollen tube ruptures and releases its enclosed sperm cells. In the last step of double fertilisation, which is characterised by defined Ca^2+^ signals in the female gametes, the sperm cells subsequently fuse with the egg cell and the central cell to form the zygote and the endosperm, respectively (Dresselhaus and Franklin-Tong, 2013; Bleckmann et al., 2014; Denninger et al., 2014; Hamamura et al., 2014; Sprunck, 2020).

To protect the male gametophyte from environmental influences on its way to a compatible flower, it is metabolically inactive, desiccated, and encapsulated in a thick and rigid pollen coat, forming the pollen grain. Once on the stigma of a compatible flower, the vegetative cell needs to be activated and polarise the tip growth machinery to a defined subcellular region to germinate from the pollen grain (Edlund et al., 2004; Rudall and Bateman, 2007). Pollen grains have apertures, areas in which the pollen coat is thinner, which predefine the possible emergence regions of the pollen tube in most angiosperms. However, in some species, such as *Arabidopsis thaliana,* the pollen emergence site is independent of these apertures and is predominantly defined by the contact site to the papilla cells. This requires sensing the contact site and a growth machinery, which can be polarised independently of the pollen morphology to loosen the pollen coat locally and allow the pollen tube’s subsequent polar emergence (Edlund et al., 2004). Moreover, in most Angiosperms, the number of pollen grains exceeds the number of female gametes, causing competition between the individual pollen grains. Thus, the rapid establishment of cell polarity and polar growth initiation required for pollen germination is crucial in this competition and is decisive for fertilisation success. (Dresselhaus et al., 2016; Sprunck, 2020). The factors responsible for sensing the papilla-pollen contact site and the polarisation of the tip growth machinery in pollen grains are unknown. Moreover, the proteins required to transmit this polarisation signal to the tip growth machinery have yet to be discovered. Therefore, we investigated this crucial aspect of pollen germination.

During pollen tube tip growth, multiple RECEPTOR-LIKE KINASES (RLKs), like POLLEN RECEPTOR KINASEs (PRKs), BUDDHA’S PAPER SEAL (BUPS), or ANXUR (ANX) RLKs were shown to be required for pollen tube growth. BUPS and ANX RLKs are crucial for maintaining pollen tube integrity and preventing the rupture of germinated pollen tubes (Boisson-Dernier et al., 2009; Miyazaki et al., 2009; Ge et al., 2017). PRKs promote general pollen tube growth and are required for chemotaxis towards the female gametophyte (Mu et al., 1994; Tang et al., 2004; Chang et al., 2013; Takeuchi and Higashiyama, 2016). PRKs were also proposed to sense stigmatic signal peptides in Tomato and thus activate pollen germination, but no general germination-promoting function for these RLKs is shown (Muschietti et al., 1998; Tang et al., 2004; Chang et al., 2013). Thus, it is still unknown whether PRKs play a general role in pollen activation and germination or which proteins are crucial for initiating pollen activation.

In various cell types and processes, RLKs activate RHO OF PLANTS (ROP) signalling pathways to establish cell polarity, promote polar growth, or confer immune responses (Kaothien et al., 2005; Duan et al., 2010; Feiguelman et al., 2018; Liu et al., 2021; Lin et al., 2022). ROP signalling pathways are mediated by plant-specific ROP GTPases, which are part of the Rho family of small GTP-binding proteins that act as molecular switches and cycle between an inactive GDP-bound state to an active GTP-bound state (Lin et al., 1996; Kost et al., 1999b; Feiguelman et al., 2018). In their active state, ROPs interact with ROP INTERACTING PARTNER (RIP) and ROP INTERACTING CRIB-CONTAINING PROTEIN (RIC) proteins, which facilitate the specific activation of downstream pathways that are required for polar growth (Holdaway-Clarke and Hepler, 2003; Shichrur and Yalovsky, 2006; Nagawa et al., 2010; Steinhorst and Kudla, 2013; Feiguelman et al., 2018). During pollen tube tip growth, ROPs are essential for cell polarisation and to promote tip growth (Lin et al., 1996; Kost et al., 1999a). Recently, it was shown that ROP signalling is additionally crucial for pollen germination, as the quadruple mutant *rop1;3;5;9* of all redundant, pollen-expressed ROPs is sterile and incapable of pollen germination (Xiang et al., 2023). As ROP signalling is required for pollen germination, we hypothesise that activators of ROP signalling are also crucial for pollen germination. However, it is unknown which ROP activators are required for pollen germination.

The activation of ROPs is stimulated by ROP-specific GUANINE EXCHANGE FACTORS (ROPGEFs), which facilitate the exchange from GDP to GTP. *Arabidopsis thaliana* has 14 of these ROPGEFs, hereafter called GEFs, which contain a conserved PLANT-SPECIFIC ROP NUCLEOTIDE EXCHANGER (PRONE) domain and variable termini (Berken et al., 2005; Bos et al., 2007; Zhang et al., 2010; Lin et al., 2012; Miyawaki and Yang, 2014). The PRONE domain forms a homodimer in which each protein binds a ROP protein and catalyses the nucleotide exchange of the GTPase, as demonstrated by the structure of ROP4 together with the PRONE domain of GEF8 (Thomas et al., 2007; Berken and Wittinghofer, 2008). Individual GEFs exhibit specific expression patterns and functions in different cell types and polarity processes. We showed that GEF3, together with the previously known GEF4, is highly expressed in root hairs and promotes polarity establishment or root hair growth (Duan et al., 2010; Denninger et al., 2019). Specific GEFs are establishing the membrane domains required for xylem development, and we showed that particular GEFs are expressed during phloem differentiation (Nagashima et al., 2018; Roszak et al., 2021). Additionally to other processes, such as hormone signalling or pavement cell morphogenesis, GEFs were extensively studied in pollen tube tip growth (Kaothien et al., 2005; Zhang and McCormick, 2007; Yu et al., 2012; Chang et al., 2013; Feiguelman et al., 2018; Lin et al., 2022).

Of the 14 GEFs of *Arabidopsis thaliana*, multiple are expressed in mature pollen grains, and four are reliably detected in transcriptomic and proteomic approaches (Supplemental Figure S1) (Mergner et al., 2020). Previous studies mainly focused on GEF12 and investigated its role in growing pollen tubes and activating ROP signalling during tip growth. This showed that the activity of GEFs is promoted by phosphorylation of their C-termini by PRK RLKs or controlled by the cytosolic AGCVIII protein kinases AGC1.5 and AGC1.7, which phosphorylate the PRONE domain of GEFs. Overexpression or misregulation of GEFs resulted in a loss of polarity, pollen tube swelling, and unidirectional growth, while loss of GEF function led to shorter pollen tubes (Zhang and McCormick, 2007; Nagawa et al., 2010; Chang et al., 2013; Zhao et al., 2013; Li et al., 2018; Li et al., 2020). As for ROPs, redundancy between multiple GEFs was indicated during pollen tube growth. A *gef1/gef9/gef12/gef14* quadruple mutant displayed a mild reduction in pollen tube length, but only two of the mutated *GEFs* are expressed in pollen (Chang et al., 2013). Recently, a *gef8/gef9/gef11/gef12/gef13* quintuple mutant of all known pollen-expressed *GEFs* showed a decrease in pollen tube integrity during tip growth and reduced fertility (Zhou et al., 2023). These studies show the importance of GEFs in promoting and maintaining polar growth and, thus, male fertility. However, they only investigated pollen tube tip growth. The role of GEFs during pollen germination and which of these GEFs initiates and activates the growth process still need to be discovered.

Compared to the redundancy found among ROPs, GEFs can have specific functions during the establishment of a new polar domain and polar growth initiation. In root hairs, we showed that establishing the polar growth domain and tip growth are two processes activated by distinct GEFs. GEF3 is required to establish the root hair initiation domain and the polarisation of ROP2, while GEF4 drives the subsequent tip growth (Denninger et al., 2019). In pollen tube germination, a polar growth domain must be established quickly from a dormant cell to provide an advantage over competing pollen. Moreover, in Arabidopsis, this polar protein accumulation must be spatially flexible, as the area of pollen tube germination is not predetermined and is defined by the region in contact with the papilla cells (Edlund et al., 2004; Dresselhaus and Franklin-Tong, 2013). Therefore, pollen germination is a great model for studying the de novo formation of polar protein domains at the plasma membrane and understanding the spatiotemporal processes required to initiate polar growth. As GEF proteins have not been investigated during pollen germination, we investigated their role in this process to understand how GEFs activate ROP signalling in a spatiotemporally controlled manner to allow the *de novo* establishment of polar cellular growth.

We focused our research on the pollen-specific ROPGEFs (GEF8, GEF9, GEF11, GEF12, and GEF13) during pollen germination (Mergner et al., 2020) and show that GEFs are distinctively crucial for pollen germination and male fertility. We demonstrate that GEF8 and GEF9, but not GEF11 and GEF12, form a transient polar domain at the plasma membrane of the pollen germination site and that these GEFs drive polar ROP activation and cellular growth. This novel subcellular localisation highlights that GEFs have specific roles during cellular processes. Furthermore, this GEF accumulation appears in a biphasic temporal manner and can shift its location. This is the first description of a polar domain with such flexibility, which is crucial for polarity establishment, hence, pollen function and fertilisation success.

## Results

### GEF8 and GEF9 biphasically accumulate at the pollen germination site

In flowering plants, pollen germination is essential for male fertility and, thus, successful double fertilisation. Therefore, it is crucial to understand the molecular mechanisms activating pollen germination upstream of the essential ROP proteins (Xiang et al., 2023). Even though multiple GEFs were shown to be involved in pollen tube growth, the GEFs involved in ROP activation during pollen germination still need to be determined (Chang et al., 2013; Zhou et al., 2023). To investigate the role of GEFs during pollen tube germination, we focused on five GEF proteins (GEF8, GEF9, GEF11, GEF12, and GEF13) that had been found to be expressed in mature pollen (Supplemental Figure S1) (Mergner et al., 2020). We fused these five GEFs N-terminally with the yellow fluorescing protein mCitrine (mCit) under the control of their endogenous promoter fragments to confirm their expression. We found mCit-GEF8, mCit-GEF9, mCit-GEF11, and mCit-GEF12 signals in mature pollen, but no signal was observed for mCit-GEF13 (Figure 1 and Supplemental Figure S2). We assessed mCit-GEF localisation by live cell imaging in germinating pollen grains *in vitro* on pollen germination medium (PGM) (Vogler et al. 2014). Shortly after imbibition on PGM, all GEFs showed a similar localisation and were evenly distributed in the cytoplasm (Figure 1, Supplemental Figure S2 and Supplemental Video 1-3). After several minutes of imbibition on PGM, we observed that mCit-GEF8 and mCit-GEF9 accumulated at a defined region in the cell periphery, which strongly correlated with the future germination site. Such an accumulation was never observed for mCit-GEF11, mCit-GEF12, or mCit-GEF13 expressed under the control of a *GEF12* promoter (Figure 1, Supplemental Figure S2, and Supplemental Video 1-3, 19-20). This indicates a specific function of GEF8 and GEF9 during pollen germination initiation. To further characterise this behaviour, we quantitatively compared the timing of protein accumulation. We defined the first frame of a visibly emerged pollen tube as a reference timepoint 0, with negative time points before and positive time points after this reference point. By quantifying the timing of mCit-GEF8 and mCit-GEF9 accumulations using multiple kymographs, we observed that both proteins showed a biphasic accumulation that, on average, slightly differed between mCit-GEF8 and mCit-GEF9 (Figure 1A-D). The initial accumulation of mCit-GEF8 and mCit-GEF9 started in a small region of the pollen grain periphery and grew to 3-4 µm within, on average, 3 minutes for mCit-GEF8 and 5 minutes for mCit-GEF9. The maximum of this initial accumulation was reached, on average, at timepoint -11 min before germination for mCit-GEF8 and -9 min for mCit-GEF9, depicting a faster and earlier accumulation of GEF8 compared to GEF9 (Figures 1C and 1D). However, it should be noted that the timing of mCit-GEF8 accumulation was more consistent, which leads to a clearer profile of the average intensity blot. In contrast, mCit-GEF9 accumulation was stronger but less consistent in its timing, which causes a less concise profile of the average intensity blot (Figures 1C and 1D). For mCit-GEF8 and mCit-GEF9, the timing of this initial accumulation, which we considered the germination initiation, was variable, and the timepoint of the maximal accumulation is specifically indicated (Figure 1A, “germination initiation”). The overall timing and persistence of this protein accumulation can be observed in the corresponding kymographs (Figure 1B). The initial accumulation of mCit-GEF8 and mCit-GEF9 did not persist at the future germination site but disappeared after, on average, 3 minutes for mCit-GEF8 and 7 minutes for mCit-GEF9. Around the time of germination, we observed a second accumulation of GEF8 and GEF9 around the initial accumulation site, which also differed in timing between both proteins. mCit-GEF8 reaccumulated at timepoint - 2 min before pollen germination, while GEF9 accumulation started after germination (Figure 1C and 1D). This slightly different timing of the biphasic accumulation indicates that during the initiation phase of pollen germination, either GEF8 or GEF9 are accumulated at the pollen germination site. Interestingly, we observed that the second accumulation of mCit-GEF8/ mCit-GEF9 was sometimes slightly shifted laterally compared to the first accumulation before germination. However, both sites were always near each other, indicating a certain flexibility during the assembly of all required proteins of the tip growth machinery (Figure 1A and 1B and Supplemental Video 1). Such flexibility of the polar growth domain is crucial for pollen function, especially in species like *Arabidopsis thaliana*, in which the pollen apertures do not predetermine the pollen germination site (Edlund et al., 2004). To confirm the accumulation and timing of GEF8 and GEF9 compared to the evenly distributed GEF12, we simultaneously observed mCit-GEF8 or mCit-GEF12 coexpressed with mScarlet-GEF9 (mSct-GEF9). mCit-GEF8 and mSct-GEF9 both accumulated with a similar timing at the same location, confirming the observations in the single marker lines. However, mCit-GEF12 was still evenly diffused in the cytoplasm, with no significant accumulation when a clear accumulation of mSct-GEF9 was visible (Figure 1E and 1F). These results show differences between individual GEFs, as mCit-GEF11, mCit-GEF12, and mCit-GEF13 do not show specific localisations during pollen germination, while mCit-GEF8 and mCit-GEF9 specifically accumulate at the pollen germination site. Moreover, mCit-GEF8 and mCit-GEF9 show a similar but distinct biphasic accumulation at the pollen germination site, with temporal differences between both proteins. Taken together, the differential localisation during pollen germination indicates a specific and subcellular localised role for GEF8 and GEF9 in initiating pollen germination.

**Figure 1:**
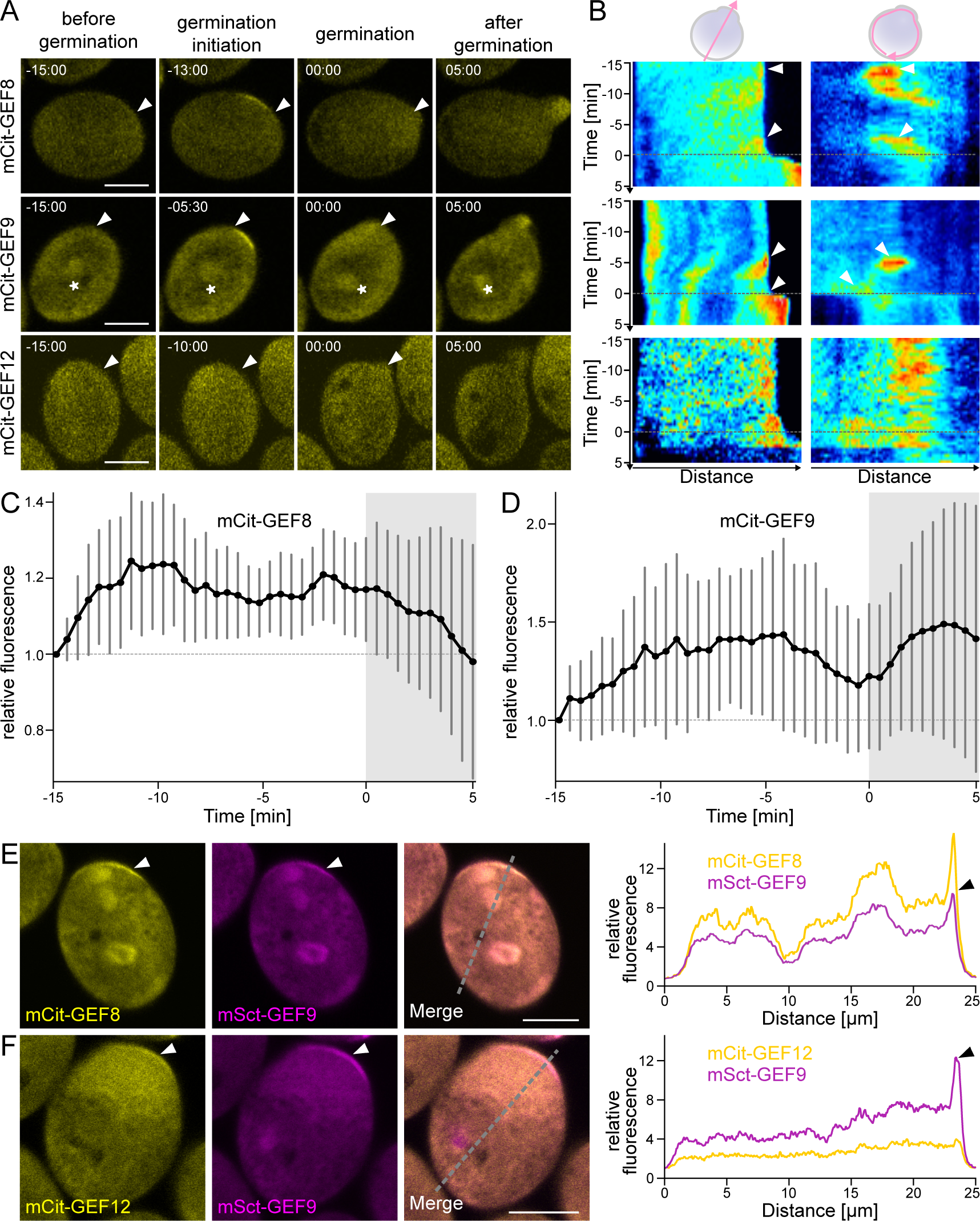
GEF8 and GEF9 specifically accumulate at the pollen germination site before germination initiation. (**A**) Protein localisation of mCit-GEF8, mCit-GEF9, and mCit-GEF12 under their respective promoters during pollen germination. Timepoint 0 corresponds to the beginning of pollen tube emergence, arrowheads mark the site of pollen emergence, and asterisks mark mCit-GEF9 localisation around the sperm cells. (**B**) Kymographs of time-lapse images corresponding to (A) along a line crossing the pollen through the germination site (left) or around the pollen grain (right). The dotted line indicates Timepoint 0 of pollen tube emergence; arrowheads highlight protein accumulations at the pollen germination site. (**C** & **D**) Average intensity profiles of mCit-GEF8 (n=13) and mCit-GEF9 (n=12) at the pollen germination site in relation to the opposite side of the pollen grain during pollen germination. (**E** & **F**) Colocalization (left) and intensity profiles along a line, as indicated in the merged image, across the pollen grain through the pollen germination site (right) of mCit-GEF8 with mSct-GEF9 (E) and mCit-GEF12 with mSct-GEF9 (F) expressed under their respective promoters. All scale bars represent 10 µm.

### Distinct GEFs are required for pollen germination *in vitro*

In light of the specific accumulation of GEFs during pollen germination, we used loss-of-function mutant lines to investigate the function of GEFs in pollen germination. We used available T-DNA lines for *gef9-t1* (*GK-717A10*)*, gef11-t1 (SALK_126725), and gef12-t1 ([SALK_103614](https://www.arabidopsis.org/servlets/TairObject?name=GK-717A10-025309&type=polyallele)*), and generated CRISPR-Cas9 full-length deletion mutants, resulting in *gef8-cΔ1, gef8-cΔ2, gef9-cΔ1, gef12-cΔ1* single mutants, and double mutants *gef8-Δc1/gef12-Δc1* and *gef8c-Δ3/gef9-cΔ2* (Supplemental Figure S3). Col-0 was used as a wild-type reference and reached a pollen germination efficiency of 84 % 4 hours after imbibition on PGM (Figure 2A-2B). *gef11-t1* (86 %) showed germination efficiencies similar to Col-0, while *gef9-t1* (79 %)*, gef12-t1* (70 %), and *gef12-cΔ1* (78 %) were slightly lower but not significantly different from Col-0. In comparison to *gef9-t1*, *gef9-cΔ1* (55 %) showed a significant reduction of pollen germination efficiency, indicating partial remaining GEF9 function in *gef9-t1*, which could be explained by the location of the T-DNA insertion in the fifth intron. We excluded the presence of full-length mRNA in *gef9-t1* by RT-PCR on flower cDNA and could only find partial *GEF9* mRNA in front and behind the T-DNA insertion site (Supplemental Figure S3). The germination efficiency of *gef8* CRISPR-Cas9 lines, *gef8-cΔ1* (63 %), and *gef8-cΔ2* (64 %) were comparable to *gef9-cΔ1* and were significantly different from Col-0. We were able to rescue *gef8-cΔ1* with the *GEF8p*::mCit-GEF8 construct, which led to a germination efficiency of 83%, proving that the absence of GEF8 causes the phenotype and confirming the functionality of the mCit-fusion constructs (Figure 2A). The pollen germination was more severely reduced in *gef8-cΔ1;9-t1* (36 %) and *gef8-cΔ3;9-cΔ2* (50 %) double mutants, while *gef8-cΔ1;12-t1* (70 %) and *gef9-t1;12-t1* (66 %) double mutants displayed no significant reduction of germination efficiency in comparison to the *gef8* and *gef9* single mutants. However, we still observed significant pollen germination in *gef8;gef9* double mutants. Therefore, we generated a *gef8-cΔ1;9-t1;12-t1* triple mutant. In this triple mutant line, *in vitro* pollen germination was almost completely abolished (0.2 %), showing that these three GEFs are essential for pollen germination (Figure 2A). In summary, this showed that GEF11 has no function in the pollen germination of *Arabidopsis thaliana*. GEF8 and GEF9 play a major role in pollen germination, as they both display a significant deficiency as single mutants, and this effect is amplified in the double mutant. The normal pollen germination efficiency of GEF12 single mutants shows a minor function of GEF12 in this process. However, because pollen germination was only abolished in the combination of *gef8*, *gef9*, and *gef12*, a partial redundancy between these three GEFs can be assumed.

**Figure 2:**
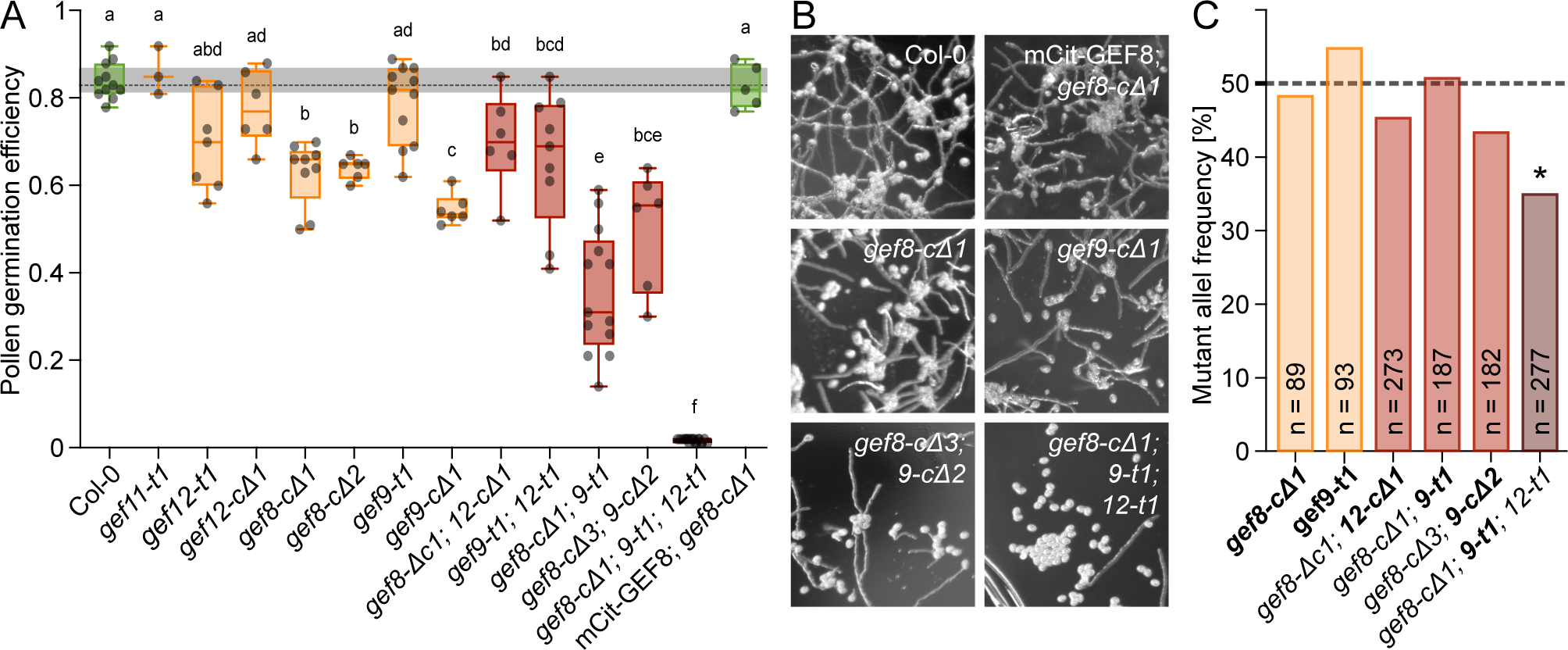
GEF8, GEF9, and GEF12 are necessary for pollen germination and male fertility. (**A**) Quantification of pollen germination efficiency of *in vitro* germinated pollen 4 h after imbibition on pollen germination media (PGM). Each point represents one replicate with more than 150 pollen grains. Tukey’s multiple comparison test was performed by one-way ANOVA, and groups of statistically significant differences are indicated with letters (*p* < 0.0001). (**B**) Representative images of *in vitro* germinated pollen 4 h after imbibition on PGM. (**C**) Quantification of mutant allele frequency in F1 generation of reciprocal crosses with Col-0 as female and the indicated mutants as pollen donors. The heterozygous allele of each genotype is indicated in bold. Asterisks indicate a significant difference in the allele frequency from the expected 50% according to Χ^2^-test (*p* < 0.05).

### GEF function is crucial for efficient male fertility *in vivo*

After we showed the necessity of GEFs for pollen germination *in vitro*, we assessed their role in pollination and male fertility *in vivo.* To determine the transmission efficiency of *gef* mutant alleles, we performed reciprocal crosses using Col-0 with *gef8, gef9, and gef12* mutant combination lines, in which one *gef* allele was heterozygous, leading to an expected transmission of 50 % for this *gef* allele in the F1 offspring (Figure 2C and Supplemental Figure S4). In the control direction with *gef* mutant lines as female, pollinated with Col-0 pollen, we did not find any significant reduction in the transmission of the mutant alleles, showing that *gef8*, *gef9*, and *gef12* are not required for female fertility (Supplemental Figure S4). When using *gef* mutant lines for pollination, we did not see significant reductions from the expected transmission *in gef8, gef9, and gef12* single and double mutant combinations. However, it needs to be mentioned that we observed nonsignificant reductions of the allele frequency in some *gef8/gef9* mutant allele combinations, which were not seen in other allele combinations of the same genes (Figure 2C and Supplemental Figure S4). Only the triple mutant *gef8-cΔ1;9-t1+/-;12-t1* had a reduced transmission of the *gef9-t1* allele (35 %), which significantly differed from the expected 50 % (p=0.0004) transmission (Figure 2C and Supplemental Figure S4). This effect on male fertility was lower than expected from the *in vitro* experiments but confirmed that *GEF8, GEF9 and GEF12* are crucial for male fertility, likely by promoting efficient pollen germination.

### The C-terminal domain is necessary and sufficient for GEF8 and GEF9 accumulation

We showed that GEF8 and GEF9 have distinct localisation compared to GEF11 and GEF12. To investigate which GEF proteins’ components are responsible for this specificity, we analysed different domains of GEFs during pollen germination. GEFs consist of a conserved catalytic PRONE domain and variable N- and C-terminal regions. The existing crystal structures of the PRONE domain of GEF8, together with ROP4, showed that these proteins form a tetramer in which two GEFs dimerise, each having one ROP bound (Thomas et al., 2007). However, the terminal regions of GEF8 were not represented in this structure, as they are thought to be intrinsically disordered. Protein structure predictions of GEF8 and ROP1 using AlphaFold2 and matched on the existing PRONE8-ROP4 structure (RCSB-PDB: 2NTY) with ChimeraX confirmed that these terminal regions are primarily unstructured, with only small helical elements (Thomas et al., 2007; Mirdita et al., 2022; Meng et al., 2023). However, in this predicted structure, both termini wrap around the protein, with the N-terminus blocking the binding site of the second GEF and the C-terminus blocking the binding site of the ROP (Figure 3A). This structure could explain the inhibitory function of both termini that was described previously (Gu et al., 2006; Zhang and McCormick, 2007). We made several mutant and deletion constructs to understand the function of GEF8 and GEF9 during pollen germination and unravel the protein features responsible for their specific biphasic accumulation (Figure 3B and 3C). We started by deleting the variable N-terminus of GEF8 (mCit-GEF8^ΔN^) and GEF9 (mCit-GEF9^ΔN^). The N-terminal deletion did not abolish the specific localisation of either protein, as we could still observe an accumulation at the germination site minutes before pollen tube emergence (Figure 3C Supplemental Video 4-5). Different effects were observed in constructs in which the variable C-terminal region of GEF8 (mCit-GEF8^ΔC^) and GEF9 (mCit-GEF9^ΔC^) was deleted. mCit-GEF8^ΔC^ and mCit-GEF9^ΔC^ were strictly cytosolic and showed no membrane attachment during pollen germination. (Figure 3C and Supplemental Video 6-7). This loss of accumulation of mCit-GEF8^ΔC^ and mCit-GEF9^ΔC^ showed that the GEF C-terminus is necessary for membrane attachment and the accumulation of GEF8 and GEF9 during pollen germination. Because the C-terminal domain is required for the accumulation of GEF8/GEF9 and this accumulation is not seen for mCit-GEF12 throughout the pollen germination process, we swapped the C-terminal domain of GEF12 with those of either GEF8 (mCit-GEF12^GEF8C^) or GEF9 (mCit-GEF12^GEF9C^) (Figure 3D and Supplemental Video 8-9). GEF12^GEF8C^ and GEF12^GEF9C^ both accumulated at the pollen germination site before pollen tube emergence, similar to the native mCit-GEF8 and mCit-GEF9, which was never observed for wild-type mCit-GEF12. This showed that the C-terminal domains of GEF8 and GEF9 are necessary for the polar accumulation of these proteins. Moreover, both C-termini are sufficient to recruit other GEFs to the cell periphery of the future pollen germination site.

**Figure 3:**
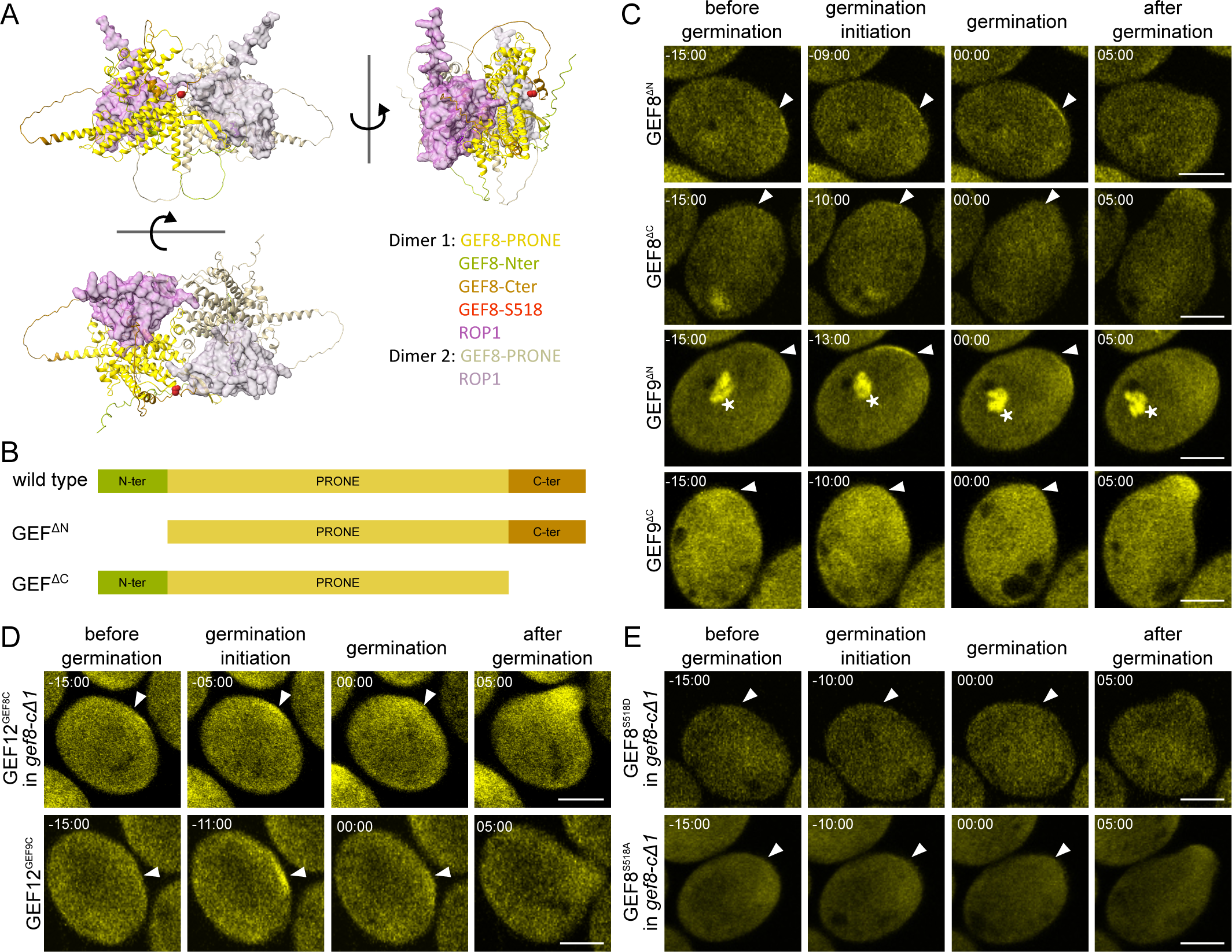
The C-terminus of GEF8 and GEF9 is necessary and sufficient for protein accumulation. (**A**) AlphaFold2 protein structure prediction of full-length GEF8 and ROP1 (Rank 1, predicted separately), matched on the PRONE8-ROP4 double-dimer structure (PDB: 2NTY). Different angles are shown, and the terminal regions are indicated with different colours. (**B**) Schemes of GEF protein structure and investigated truncation constructs, with variable N and C terminal domains indicated as in (A). (**C** – **E**) Protein localisation of different mutant constructs under their respective promoters during pollen germination. Timepoint 0 corresponds to the beginning of pollen tube emergence, arrowheads mark the site of pollen emergence, and asterisks mark localisation around the sperm cells. (**C**) Truncation constructs mCit-GEF8^ΔN^, mCit-GEF8^ΔC^, mCit-GEF9^ΔN^, and mCit-GEF9^ΔC^ in Col-0. (**D**) Domain swap constructs of GEF12 with alternative C-terminal domain, mCit-GEF12^GEF8C^ and mCit-GEF12^GEF9C^ in Col-0. (**E**) Phosphorylation site mutations of GEF8-S518 to a phospho-mimic (mCit-GEF8^S518D^) and phospho-dead variant (mCit-GEF8^S518A^) variant expressed in *gef8-cΔ1*. All scale bars represent 10µm.

### A phosphorylation site in the C-terminus of GEFs influences their accumulation

To further understand the mechanism of GEF activation and localisation, we investigated the role of phosphorylation in GEF8 and GEF9 C-terminal domains. The C-terminus of GEFs was shown to be phosphorylated by RLKs at a serine of the conserved SPxxRH motif, leading to the activation of the PRONE domain (Kaothien et al., 2005; Zhang and McCormick, 2007; Cheung and Wu, 2008). The phosphorylation at this site was recently confirmed by proteomic approaches (Mergner et al., 2020). In the predicted structure, this phosphorylation site is accessible and could affect the blocking function of this domain (Figure 3E). To elucidate any potential functional effect of the phosphorylation of this serine on pollen germination, we mutated this serine to a phospho-dead (S518A) and potentially phospho-mimic (S518D) versions of GEF8 and transformed them into the *gef8-cΔ1* background. Both mutants, GEF8^S518D^ and GEF8^S518A^, displayed no significant accumulation at the pollen tube germination site (Figure 3E and Supplemental Video 10-11). This indicates that the protein accumulation is independent of the activity status of GEF8, but rather, the phosphorylation reaction by RLKs is critical for GEF accumulation. In summary, the deletion constructs of GEF8 and GEF9, in combination with the domain swap experiments, show that the C-terminal domain is necessary for the distinct accumulation of GEF8 and GEF9 and sufficient to transfer this function onto GEF12, which usually does not accumulate in this system. In addition, GEF8^S518^ seems to have an essential role in GEF8 accumulation at the germination site, indicating that the activation by RLKs is a crucial factor in this process.

### GEF8 is necessary for ROP activation

GEFs activate ROPs by exchanging GDP for GTP, leading to the recruitment of RIP or RIC proteins and the subsequent activation of downstream processes (Feiguelman et al., 2018). For example, RIC3 and RIC4 were shown to regulate pollen growth by mediating the modulation of Ca^2+^ fluxes and F-actin formation (Theos et al., 2005). RICs contain a CDC42/RAC INTERACTIVE BINDING (CRIB) motif, which mediates the interaction with active GTP-bound ROPs. ROP1, ROP3, and ROP5 are specifically expressed in pollen grains. We observed mCit-ROP1, mCit-ROP3, and mCit-ROP5 under their endogenous promoters, and all three ROPs localised to the cytoplasm without any specific accumulation at the site of germination, as observed for mCit-GEF8 and mCit-GEF9 (Figure 4A and 4B and Supplemental Video 12-14). Nevertheless, as the CRIB domain is sufficient for active ROP binding, it allows the utilisation of the CRIB domain as a biosensor for the localisation of active ROPs (Luo et al., 2017). Here, we used the CRIB motif of RIC4 (CRIB4) fused to mCit under the control of the *GEF12* promoter fragment (*GEF12p*::CRIB4-mCit) as an indicator of ROP activity during pollen germination (Figure 4C and 4D). We observe a persistent CRIB4-mCit accumulation at the germination site during the germination process, which differs from the transient biphasic accumulation of mCit-GEF8 and mCit-GEF9 (Figure 4 and Supplemental Video 15). However, the timing of the initiation of CRIB4-mCit accumulation was similar to the one observed for both GEFs, approximately 10 minutes before germination. In addition, the overall width of the accumulation of CRIB4-mCit was similar to that of mCit-GEF8 and mCit-GEF9 (3-4 µm). To show the causality between the accumulation of GEF8 and CRIB4, we investigated CRIB4-mCit localisation in a *gef8-cΔ1* background. In *gef8-cΔ1,* CRIB4-mCit accumulation was lost, and its localisation is cytoplasmic, with no apparent accumulation at the germination site during the pollen germination process (Figure 4C and Supplemental Video 16). These results suggest that CRIB4 (a biosensor for active ROPs) accumulates at the germination site during the germination process, and this accumulation requires the activity of GEF8, which shows a functional link between GEF8/9 accumulation and ROP activation.

**Figure 4:**
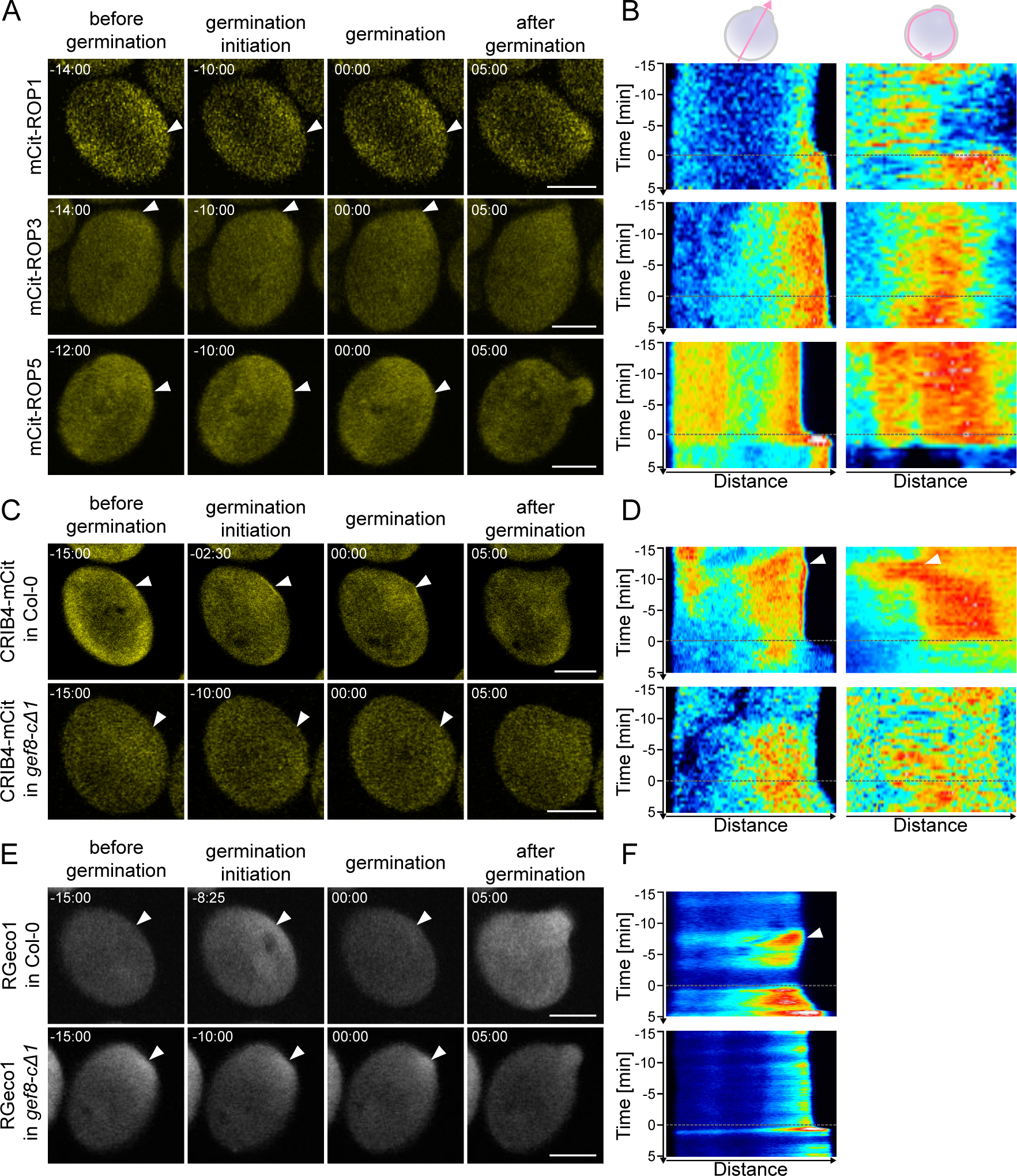
GEF8 is required for polar ROP activation and Ca^2+^ signalling. (**A**) Localization of mCit-ROP1, mCit-ROP3, and mCit-ROP5 under their respective promoter during pollen germination. Timepoint 0 corresponds to the beginning of pollen tube emergence, and arrowheads mark the site of pollen emergence. (**B**) Kymographs of time-laps images corresponding to (A) along a line crossing the pollen through the germination site (left) or around the pollen grain (right). The dotted line indicates timepoint 0 of pollen tube emergence; arrowheads highlight protein accumulations at the pollen germination site. (**C**) Localisation of the ROP activity indicator *GEF12p*::CRIB4-mCit in Col-0 and *gef8-cΔ1* backgrounds. (**D**) Kymographs of time-laps images corresponding to (C) along a line crossing the pollen through the germination site (left) or around the pollen grain (right). Arrowheads highlight protein accumulations at the pollen germination site. (**E**) *LAT52p*::RGeco1 Ca^2+^ biosensor in Col-0 and *gef8-cΔ1* background. (**F**) Kymographs of time-laps images corresponding to (E) along a line crossing the pollen through the germination site. Arrowhead highlights signal increase at the pollen germination site. All scale bars represent 10 µm.

### GEF8 oscillation leads to changes in calcium oscillation

ROP signalling leads to the activation of downstream pathways required for pollen tube growth, such as Ca^2+^ oscillations, which are essential for pollen tube growth and guidance (Cheung and Wu, 2008; Dresselhaus et al., 2016; Gao et al., 2016). Ca^2+^ undergoes fine-tuned oscillations and concentration variations to regulate polar growth (Iwano et al., 2009). Ca^2+^ fluctuations are described during pollen germination *in vivo*, but their timing to other activators of pollen germination and their cause during pollen activation still need to be understood (Iwano et al., 2004). We used the Ca^2+^-indicator RGeco1 under the control of the *Lat52* promoter to monitor Ca^2+^ oscillations during pollen germination in Col-0 (Figure 4E and Supplemental Video 17). Compared to the regular short oscillations during pollen tube growth, we observed longer and persisting Ca^2+^ elevations. We found a first increase of Ca^2+^ around - 8 minutes before germination, which persisted with different intensities for multiple minutes, decreased shortly before germination and increased again at timepoint 0 (Figure 4E and 4F). This initial increase of Ca^2+^ was comparable to mCit-GEF8 and mCit-GEF9 accumulation, indicating a possible interaction between GEF8/9 and Ca^2+^ elevations. To investigate the association between GEF function and Ca^2+^ oscillation, we observed RGeco1 in *gef8-cΔ1* during pollen germination (Figure 4E and Supplemental Video 18). Interestingly, the Ca^2+^ elevation pattern was not abolished but differed from the one observed in Col-0. Compared to Col-0, the RGeco1 signal did not increase in a persistent Ca^2+^ elevation in the *gef8-cΔ1* mutant but rather showed several lower intensity peaks throughout the germination process, with one stronger Ca^2+^ increase at the germination timepoint (Figure 4E and 4-F). This result further emphasises the necessity of GEF8 for ROP activation, resulting in normal Ca^2+^ elevations and pollen germination. In summary, the absence of GEF8 impacted CRIB4-mCit (active ROP biosensor) and Ca^2+^ oscillations, showing the crucial function of GEFs in activating ROP signalling leading to pollen germination.

## Discussion

Pollen germination is a critical step in plant fertilisation, and characterising the underlying protein functions is crucial for understanding the activation of this process. Here, we identified specific ROPGEFs required for pollen germination and male fertility. These GEFs have distinct functions during pollen germination, which are conferred, in parts, by their C-terminal domain. Five of the 14 GEFs are expressed in mature Arabidopsis pollen (Supplemental Figure S1). We confirmed the presence of GEF8, GEF9, GEF11, and GEF12 using translational fusion lines but not GEF13 (Figure 1 and Supplemental Figure S2). As recent proteomics analysis only found very low amounts of GEF13 in mature pollen, it remains unclear whether GEF13 is irrelevant for pollen tube growth or is very specifically translated (Mergner et al., 2020). Previous results also indicated the presence of GEF1 and GEF14 during pollen tube growth, but the detection of the expression level of these two genes was found to be low and inconsistent (Gu et al., 2006; Zhang and McCormick, 2007; Chang et al., 2013). In line with this, *GEF1* and *GEF14* were not identified in recent transcriptomics or proteomics data (Mergner et al., 2020).

The four consistently expressed GEFs have very distinct localisations, with GEF11 and GEF12 evenly distributed in the cytoplasm throughout the germination process, while GEF8 and GEF9 accumulated at the site of pollen germination before pollen tube growth (Figure 1 and Supplemental Figure S2). A similar accumulation of GEFs was described for GEF3 and GEF14 during root hair initiation (Denninger et al., 2019). However, during root hair initiation, the polar domain requires around 30 minutes to be established, is locally fixed and is persistent for hours (Denninger et al., 2019). The protein accumulations we observed for GEF8 and GEF9 were established significantly faster but shorter, as they only persisted at the pollen germination site for several minutes (Figure 1). An exciting aspect and a difference to the initiation of root hair growth is that the localisation of GEF8 and GEF9 accumulation was not fixed but could shift laterally during the initiation of pollen germination. This could be an essential feature of polar growth in pollen, as these cells need to adjust the germination site in response to contact with papilla cells. Thus, the flexibility of this growth machinery is required to initiate growth at the optimal position in species such as *Arabidopsis thaliana* with no predetermined germination site at the pollen apertures (Edlund et al., 2004). We also found differences in the timing of protein accumulation between GEF8 and GEF9, leading to a constant presence of GEF8 or GEF9 at the pollen germination site but with different ratios to each other (Figure 1). It remains to be determined whether they have redundant functions at this location or if distinct pathways are activated by these particular proteins, as described for downstream RIC proteins (Gu et al., 2005).

The *gef8* and *gef9* single mutants had significantly reduced pollen germination efficiencies, while *gef11* and *gef12* single mutants did not display significant reductions in pollen germination (Figure 2 and Supplemental Figure S3). Compared to the previously postulated redundancy of GEFs during pollen germination and growth, our results suggest, at least during pollen germination, that individual GEFs have distinct localisations and non-redundant functions (Chang et al., 2013; Zhou et al., 2023). The availability of suitable mutant lines can explain the discrepancy with the previous reports, which did not detect the same defects, as the observed phenotypes of *gef8* and *gef9* mutants were only evident in the CRISPR-Cas9 deletion mutants generated in this study (Figure 2 and Supplemental Figure S3). The only available T-DNA line for *GEF8* had multiple T-DNA integrations and, in our experience, showed developmental defects independent of the integration in *GEF8*. The available T-DNA line of *GEF9* showed a mild and less consistent phenotype than the generated CRISPR-Cas9 deletion mutant (Figure 2 and Supplemental Figure S3). In combination with the presented localisation data, our phenotyping analysis shows that not all GEFs act redundantly during pollen germination. We suggest that these GEFs activate different aspects required for polar growth to another degree. Thus, all GEFs are distinctively required for efficient pollen germination and male fertility, which only leads to a severe phenotype in higher-order mutants (Figure 2), similar to the observations made in loss-of-function mutants of all pollen-expressed *ROPs* (Xiang et al., 2023). Additionally, the severity of the *gef8-cΔ1;9-t1;12-t1* triple mutant phenotype observed *in vitro* can be overcome *in vivo*, as the *gef8-cΔ1;9-t1;12-t1* triple mutant, which hardly showed pollen germination on PGM, still had a significant transmission in reciprocal crosses (Figure 2C and Supplemental Figure S4), which is similar to recent data shown in *gef8;9;11;12;13* quintuple mutants. Potentially, other GEFs might have a higher activity *in vivo* that could rescue the lack of these primary GEFs, leading to a weaker phenotype than the total loss of ROP activity (Xiang et al., 2023).

The distinct localisations we observed for GEF8 and GEF9 indicate that a specific feature of those proteins transmits this function. As the PRONE domain of all GEFs is conserved, we focused on the variable termini with no apparent structure and seeming intrinsically disordered (Figure 3) (Berken et al., 2005). The predicted full-length structure of GEF8 suggests that both termini are folding around the PRONE domain to either block ROP binding or prevent protein dimerisation, indicating inhibitory functions. However, the confidence in the structure of these regions is very low, and it remains unclear how these termini are folded or if they can take different shapes depending on the situation. The N-terminus was shown to have activating functions for GEF activity, while the C-terminus inhibits PRONE activity (Gu et al., 2006; Zhang and McCormick, 2007). However, in contrast to the suggested function of the terminal domains for GEF activity, we observed other functions for their localisation. The deletion of the N-terminus did not significantly alter protein accumulation behaviour. In contrast, the deletion of the C-terminus abolished protein accumulation (Figure 3C), which shows that the contribution of the termini to protein activity and localisation independently regulates different aspects of GEFs. We speculate that the C-terminus of GEF8 and GEF9 is required for interaction with RLKs like PRKs, as shown before for GEF12, as mutations in the conserved phosphorylation site can alter this accumulation (Figure 3) (Kaothien et al., 2005; Chang et al., 2013). However, this interaction must be specific to GEF8 and GEF9 as we do not see such accumulations for other GEFs. Moreover, the ability for protein accumulation before pollen tube emergence can be transferred onto GEF12 when exchanging the C-terminal domain (Figure 3D). Suggesting that other scaffolding proteins or other RLKs than the previously reported PRK2, PRK6, or BUPS are responsible for this interaction, as they also interact with GEF12 (Chang et al., 2013; Zhao et al., 2013; Takeuchi and Higashiyama, 2016; Yu et al., 2018; Zhou et al., 2021). However, it is also possible that additional mechanisms are crucial for this accumulation or that further factors transmit a specific interaction of GEF8 and GEF9 with the known RLKs at this particular time during the initiation of pollen germination. Other proteins known to show similar accumulation at the pollen germination site before germination are the ROP effector scaffold proteins RIP1, RIC1, and potentially other members of these families or BOUNDARY OF ROP DOMAIN (BDR) proteins. However, it is unlikely that these scaffolds drive GEF accumulation, as they are ROP effectors, and their accumulation depends on ROP activation (Li et al., 2008; Zhou et al., 2015; Sugiyama et al., 2019; Xiang et al., 2023). Still, these effectors may have a role in stabilising the GEF accumulation in a feedback loop to confine the domain, as shown in root hairs (Denninger et al., 2019).

The accumulation of GEFs did not lead to any significant accumulation of ROPs at the pollen germination site (Figure 4A), which conforms with recent observations of ROPs during pollen germination (Xiang et al., 2023). Still, we could observe that a marker for active ROPs and Ca^2+^ signals are elevated at the pollen germination site with a timing similar to the observed GEF accumulations. In line with this, the accumulation of the active ROP marker was lost, and the Ca^2+^ elevation pattern was altered in the *gef8-cΔ1* mutant (Figure 4C and 4D). The complete loss of accumulation of the ROP activity indicator (CRIB domain) is surprising, considering the mild phenotype in *gef8-cΔ* mutants (Figure 2). However, the accumulation of this indicator is also very low, and it might be that slight changes in ROP activity already have significant effects on this sensor, even though the remaining ROP activity is sufficient to trigger altered Ca^2+^ signals and induce pollen germination. However, this alteration of ROP activity in *gef* mutants indicates that GEF accumulation is upstream of ROP activity and requires a different mechanism. A connection to phospholipid signalling is possible, as shown in root cells (Platre et al., 2019). Candidate proteins are PHOSPHATIDYLINOSITOL 4-PHOSPHATE 5-KINASEs (PIP4Ks) that regulate phospholipid abundance and regulate tip growth in pollen tubes and root hair cells (Kusano et al., 2008; Kato et al., 2024). PIP4Ks also accumulate at the pollen germination site and might act together with GEFs to drive ROP signalling, but results in root hairs suggest that PIP4K accumulation is downstream of GEF accumulation (Denninger et al., 2019; Kato et al., 2024). Still, studying the connection to other pathways and their regulatory connection will be a challenging task to fully understand polarity establishment and polar growth initiation.

We show that specific GEFs are required for efficient pollen germination and that GEF8 and GEF9 display distinct localisations compared to GEF11 and GEF12. Together, this shows that GEFs are not redundant during pollen germination and can have specific functions within the same process in one cell. The novel polar domain of accumulated GEF8/9 protein in germinating pollen tubes was spatiotemporally flexible and not static as in previously described processes. We hypothesise that this flexibility is crucial to define the pollen germination site independently of predeterminant features and shows the *de novo* assembly of a polar growth domain.

## Material and Methods

### Plant material and growth conditions

*Arabidopsis thaliana* plants were grown on soil under long-day conditions (16 h of light at 21 C) in a growth chamber. *Arabidopsis thaliana* ecotype Col-0 was used as a wild-type reference. The mutant lines *gef9-t1* (GK-717A10), *gef11-t1* (SALK_126725C), and *gef12-t1* (SALK_103614) were obtained from NASC (Nottingham Arabidopsis Stock Centre). Single t-DNA insertion was confirmed by segregation analysis, and the t-DNA insertion site was checked by sequencing the genotyping PCR product on both sides of the insertion site. Primers used for genotyping are listed in Supplemental Table S1. Double and triple mutants were made by crossing these lines and were selected by PCR. Fluorescently labelled *GEF12p*::mCit-GEF12 was used from (Denninger et al., 2019).

### CRISPR/Cas9 deletion lines

To generate *gef8-cΔ1* and *gef12-cΔ1* CRISPR/Cas9 deletion lines, we used an egg cell-specific promoter-driven CRISPR/Cas9 (Wang et al., 2015). We used two gRNAs, one in 5’ and the other in 3’ of the gene, cloned in tandem into one vector (Supplemental Figure S3). The selection of positive transformants in T1 generation was done on ½ MS medium containing Hygromycin (20 µg/ml).

To generate CRISPR/Cas9 deletion lines for *gef8-cΔ2* (2.2kb) and *gef9-cΔ1* (2.7kb) single mutants and *gef8c-Δ3/gef9-cΔ2* double mutants, we used a multiplex editing approach based on an optimised zCas9i (Stuttmann et al., 2021). We used four gRNAs per gene, two in 5’ and two in 3’ of the gene. (Supplemental Figure S3). The selection of positive lines in T1 was performed using seed fluorescence and Basta resistance. T2 selection was made by selecting non-glowing seeds.gRNAs were determined using ChopChop (https://chopchop.cbu.uib.no/) (Labun et al., 2019) and CCTop (https://cctop.cos.uni-heidelberg.de/) (Stemmer et al., 2015) and successful cloning was confirmed by sequencing. Genotyping was performed by primers 200-300bp outside the deleted area. The characterisation of the deletion was done by sequencing the resulting PCR product. Confirmation of homozygous mutant alleles was done by combination with a primer binding in the deleted region and was confirmed in the following generation. All primers are listed in Supplemental Table S1.

### Plasmid construct

All non-CRISPR/Cas9 constructs were generated using the GreenGate cloning system (Lampropoulos et al., 2013) with modified cloning procedures. A list of combined modules and their sources is provided in Supplemental Table S2. As native promoter sequences upstream regions of the START codon, including the 5’UTRs, were cloned for *GEF8* (AT3G24620, - 2501bp), *GEF9* (AT4G13240, -1463bp), *GEF11* (AT1G52240, -973bp), *GEF12* (AT1G79860, 701bp), and *GEF13* (AT3G16130, 850bp and 1200pb) The *LAT52* promoter was amplified from LHR lines (Denninger et al., 2014) and a RGeco1 module was kindly provided by Rainer Waadt (Waadt et al., 2017) The CRIB4 module was generated by amplification of the CRIB domain of RIC4 (AT5G16490, amino acid I64-I131) as described before (Luo et al., 2017) from flower cDNA. All primers used to generate new Entry-Vector modules are listed in Supplemental Table S1. The correct amplification and cloning of entry-vector modules were confirmed by sequencing. We generated a new GreenGate-compatible destination vector to achieve a higher plant transformation efficiency. For this, we amplified the vector backbone of pHEE401E (Wang et al., 2015) and the GreenGate cloning cassette of pGGZ003 (Lampropoulos et al., 2013) by PCR (Primers in Supplemental Table S1) and combined both fragments using added MluI and BamHI sites. The resulting plasmid was confirmed by sequencing and named pGGX000.

### Phenotyping of pollen germination efficiency

For phenotyping of *in vitro* pollen germination efficiency, pollen of freshly opened flowers was germinated at 22°C on solid pollen germination medium (PGM), containing 1.5% agarose (w/v), 18% sucrose (w/v), 0.01% boric acid (w/v), 1mM CaCl_2_, 1mM Ca(NO3)_2_, 1mM MgSO_4_, 10µM 24-epibrassinolides (MedChemExpress, HY-N084824, dissolved to 5mM in Ethanol), and adjusted to pH=7.0 using 100 mM KOH,) (Vogler *et al*., 2014). Germination was analysed 4 hours after imbibition on the PGM, using a Leica MZ16 stereomicroscope equipped with DMC5400. The images were analysed using the multipoint tool of ImageJ (https://imagej.nih.gov/ij/docs/guide/146-19.html#sec:Multi-point-Tool), then making the ratio of germinated and ungerminated pollen to get the germination efficiency. Each line was analysed in at least three independent experiments, including three replicates in each experiment. Each data point represents an independent replicate analysing more than 150 pollen grains. Statistical analysis was performed using GraphPad Prism 9. We used a one-way ANOVA with a post-Tukey test (significance level <0.0001) to test for significant differences in pollen germination efficiency.

### Fluorescence imaging and quantification

Imaging of *in vitro* pollen germination was done at 22°C on solid PGM, containing 1.5% agarose (w/v, 10% sucrose (w/v), 0.01% boric acid (w/v), 5mM CaCl_2_, 5mM KCl, 1mM MgSO_4_, adjusted to pH=7.5 using 100 mM KOH (Boavida and McCormick, 2007) and supplemented with 10µM 24-epibrassinolides (MedChemExpress, HY-N084824, dissolved to 5mM in Ethanol) (Vogler et al., 2014).

Live cell imaging was performed using an Olympus confocal FV-1000 equipped with a 40x water immersion (1.15 NA) objective, an Argon laser, and a 559 nm diode laser. Signals were detected with high-sensitivity detectors. mCitrine (YFP) was excited at 515 nm and detected between 520-550nm. mScarlet (RFP) and RGeco signals were exited at 559 nm and were detected between 580-653 nm. The pinhole was set to 1AU, and images were taken with a 2x line average in a 1024x1024 pixel scanning field. The same settings were applied to all images, and the excitation laser intensity was set to a minimum for the individual lines to avoid phototoxicity. GEF live pollen germination images were acquired every 30 seconds for 30 minutes. For Lat52p::RGeco, images were acquired every 5 seconds for 30 minutes. The images were processed and analysed using ImageJ. Kymographs were generated with the MultipleKymograph plugin, using a line width of five pixels.

## Acknowledgements

We are grateful to Andrea Bleckmann, Claus Schwechheimer (Technical University of Munich), and Guido Grossmann (University Düsseldorf) for helpful comments and suggestions on our manuscript. We thank Thomas Dresselhaus (University Regensburg) for the possibility of confirming our results on an independent microscope setup and Andrea Bleckmann for imaging advice and microscope instructions. We thank all members of the Schwechheimer group for their support, especially Franziska Anzenberger, for her dedication and support of our plant work and experimental procedures. The Deutsche Forschungsgemeinschaft supported this work within the Collaborative Research Centre SFB924/3 (#170483403) #A15.

## Author contributions

P.D. conceived the study; A.M.B. and P.D. performed experiments, analysed data, prepared figures, and wrote the manuscript with input from A.L.; A.M.B., A.L. and P.D. generated experimental material; All authors read and approved the final version of the manuscript.

## Supplemental Figures

**Supplemental Figure S1:**
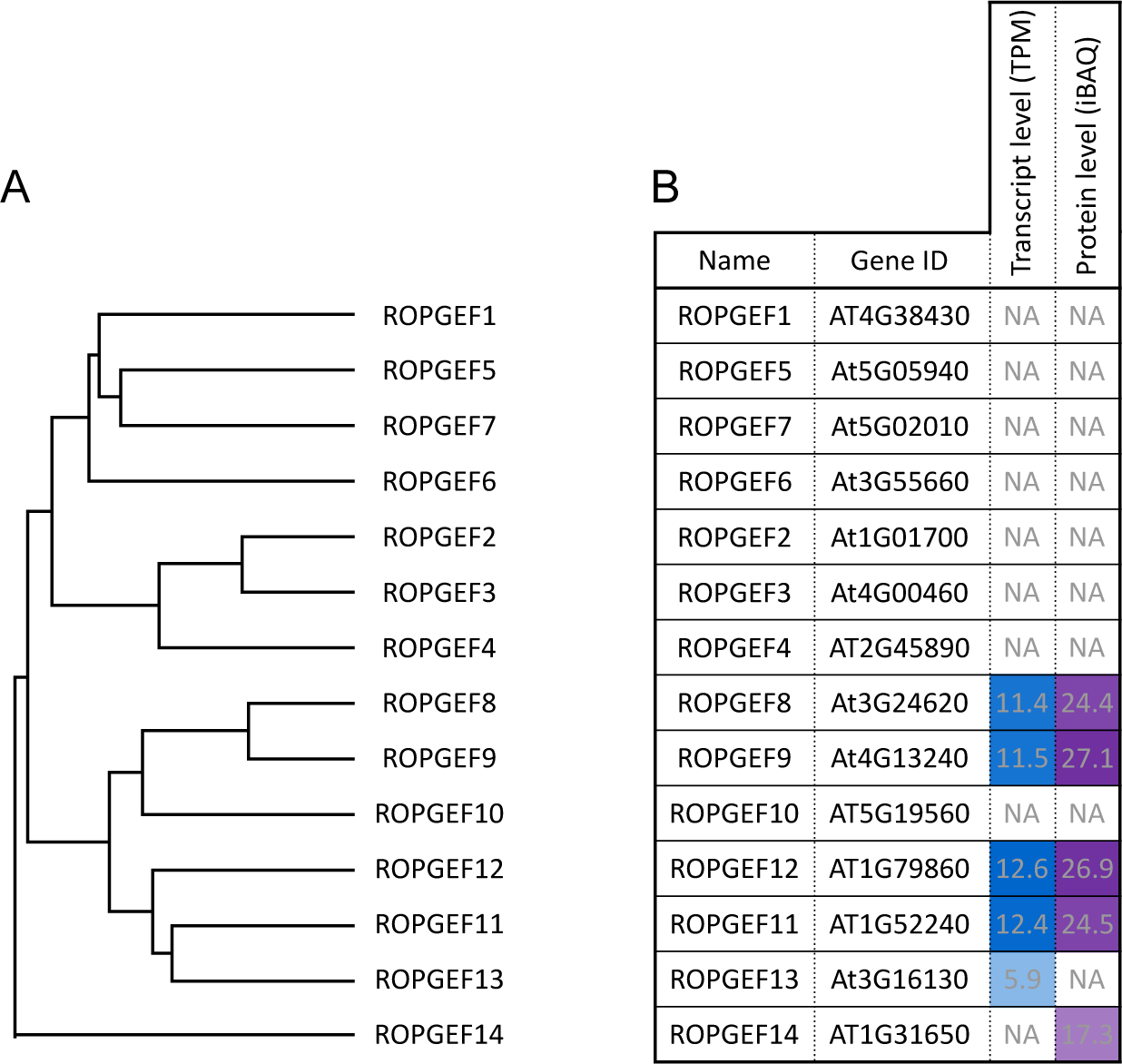
Phylogeny of all ROPGEFs of Arabidopsis thaliana and expression levels in mature pollen. (**A**) Phylogenetic tree of the 14 ROPGEFs of *Arabidopsis thaliana* after alignment of the full-length protein sequence in Jalview, using the integrated MUSCLE alignment tool and calculation of an average distance (BLOSUM62) tree. (**B**) Presence of transcript or protein for all 14 ROPGEFs in mature pollen, according to the ATHENA – Arabidopsis THaliana ExpressioN Atlas (Mergner et al., 2020). GEFs are sorted according to the phylogenetic tree. Levels of the transcript are shown in transcripts per kilobase million (TPM) and intensity-based absolute quantifications (iBAQ) for protein levels. NA indicates that no transcript or protein was detected in this tissue.

**Supplemental Figure S2:**
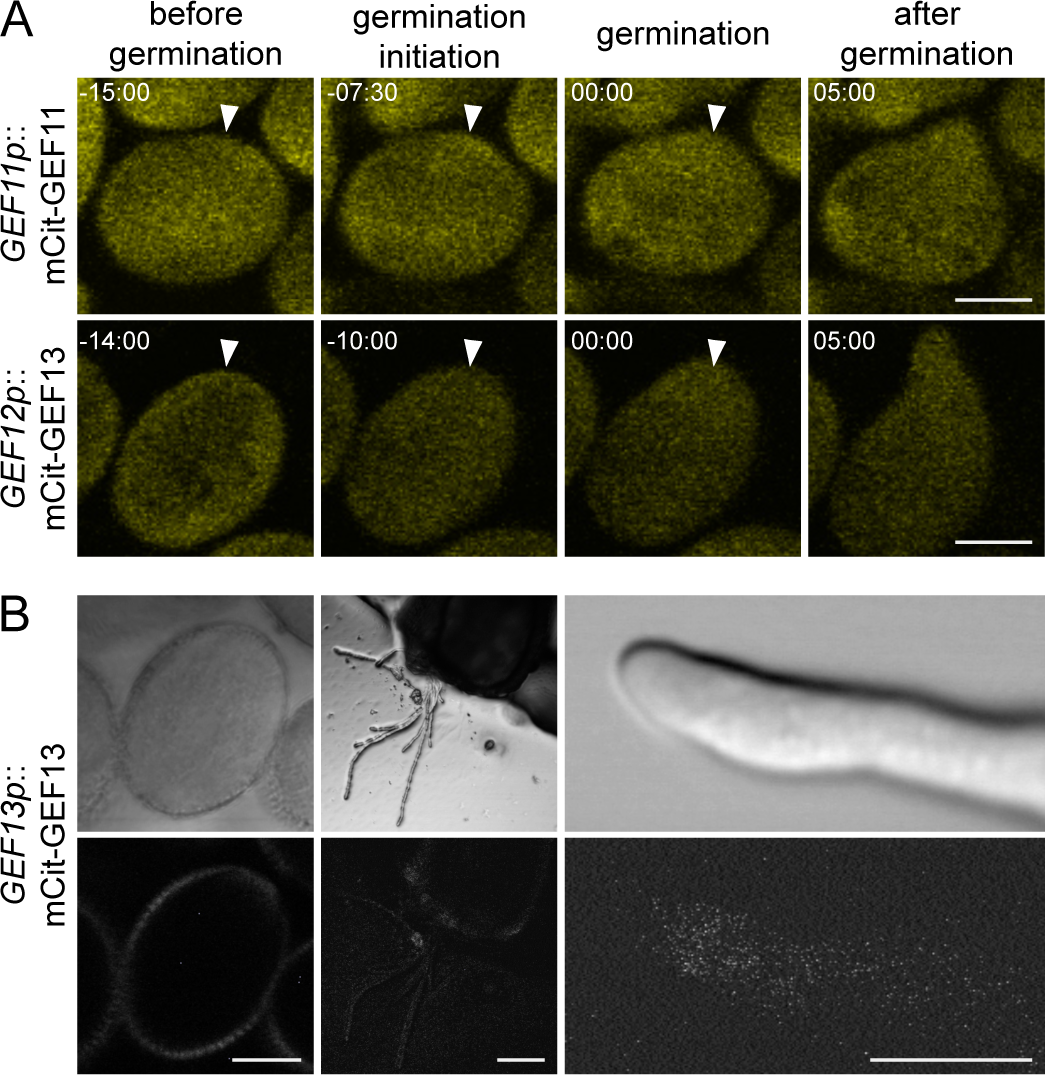
GEF11 and GEF13 do not accumulate at the pollen germination site. (**A**) Protein localisation of mCit-GEF11 under its own promoter and mCit-GEF13 under control of a *GEF12* promoter fragment during pollen germination. Timepoint 0 corresponds to the beginning of pollen tube emergence, and arrowheads mark the site of pollen emergence. (**B**) Example images of mCit-GEF13 under the control of a *GEF13* promoter fragment in mature pollen grains and pollen tubes grown through a cut pistil. In neither case, any signal could be detected. All scale bars represent 10µm.

**Supplemental Figure S3:**
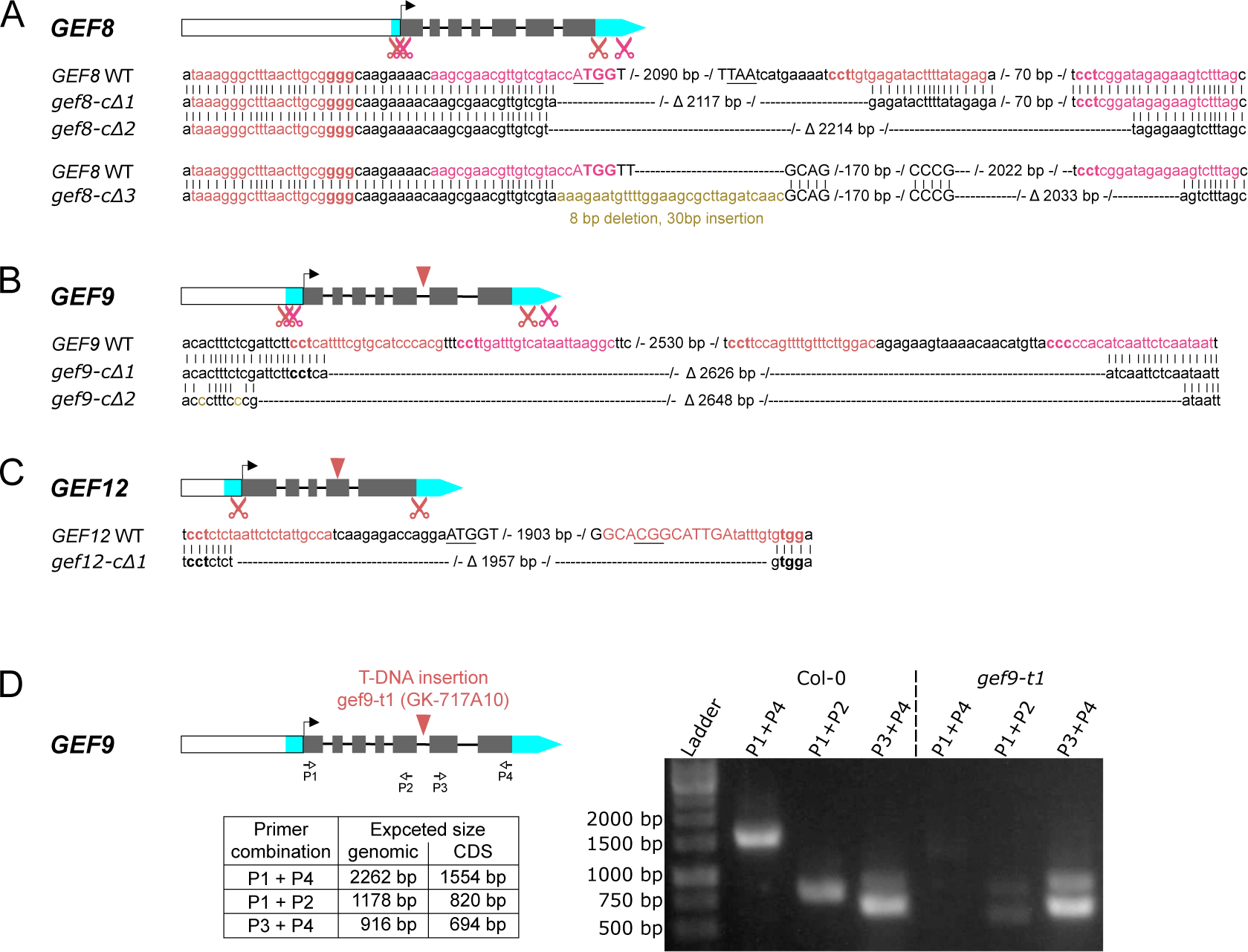
Genomic structure and mutant allele information of GEF8, GEF9, and GEF12. (**A-C**) *GEF8*, *GEF9*, and *GEF12* genomic structures with the corresponding gRNA sites (scissors) and T-DNA insertion sites (arrowhead) for *gef9-t1* (GK-717A10) and *gef12-t1* (SALK_103614). Promoter regions are shown as white boxes, UTRs in cyan, and exons in grey boxes. WT sequence for each gene and corresponding CRISPR/Cas9 deletion line is shown, and the size of CRISPR/Cas9 induced deletions is indicated. The gRNA target sequence is highlighted in colour with PAM in bold. The START and STOP codons are underlined. (**D**) *GEF9* genomic structure with T-DNA insertion sites (arrowhead) for *gef9-t1* (GK-717A10) and the location of primers used to test the mRNA presence is indicated. The table shows the expected PCR product size with the indicated primer combination for non-spliced templates (genomic) and correctly spliced templates (CDS). The gel image shows PCR products of PCRs using the indicated primer combination on cDNA from Arabidopsis flowers of Col-0 or *gef9-t1* plants.

**Supplemental Figure S4:**
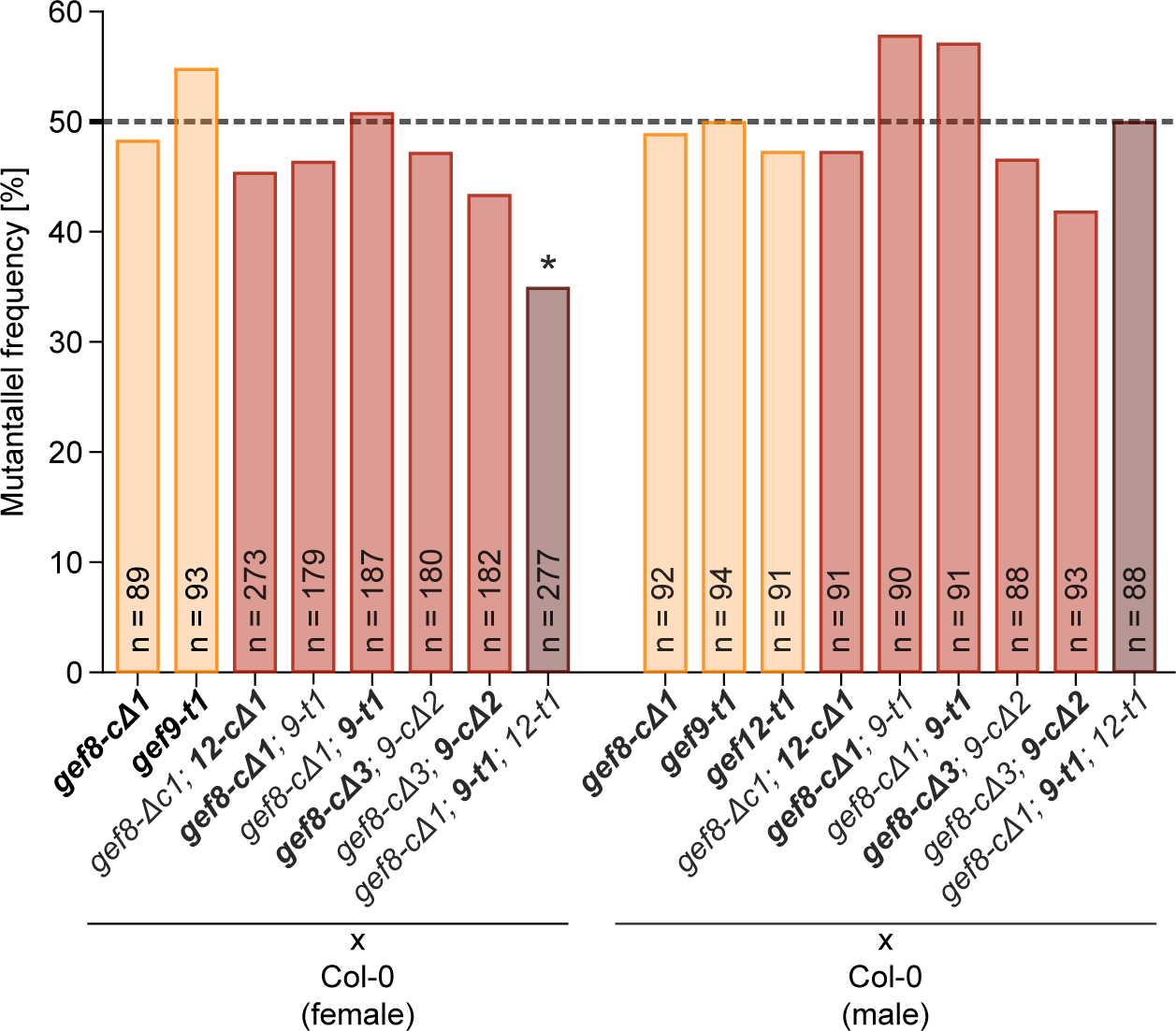
Reciprocal crosses of *GEF* mutant lines. Quantification of mutant allele frequency in F1 generation of reciprocal crosses with Col-0 as female and the indicated mutants as pollen donors (left) or indicated mutants as female and Col-0 as pollen donor (right). The heterozygous allele of each genotype is shown in bold. Asterisks indicate a significant difference in the allele frequency from the expected 50% according to Χ^2^-test (*p* < 0.05)

**Supplemental Table S1:**
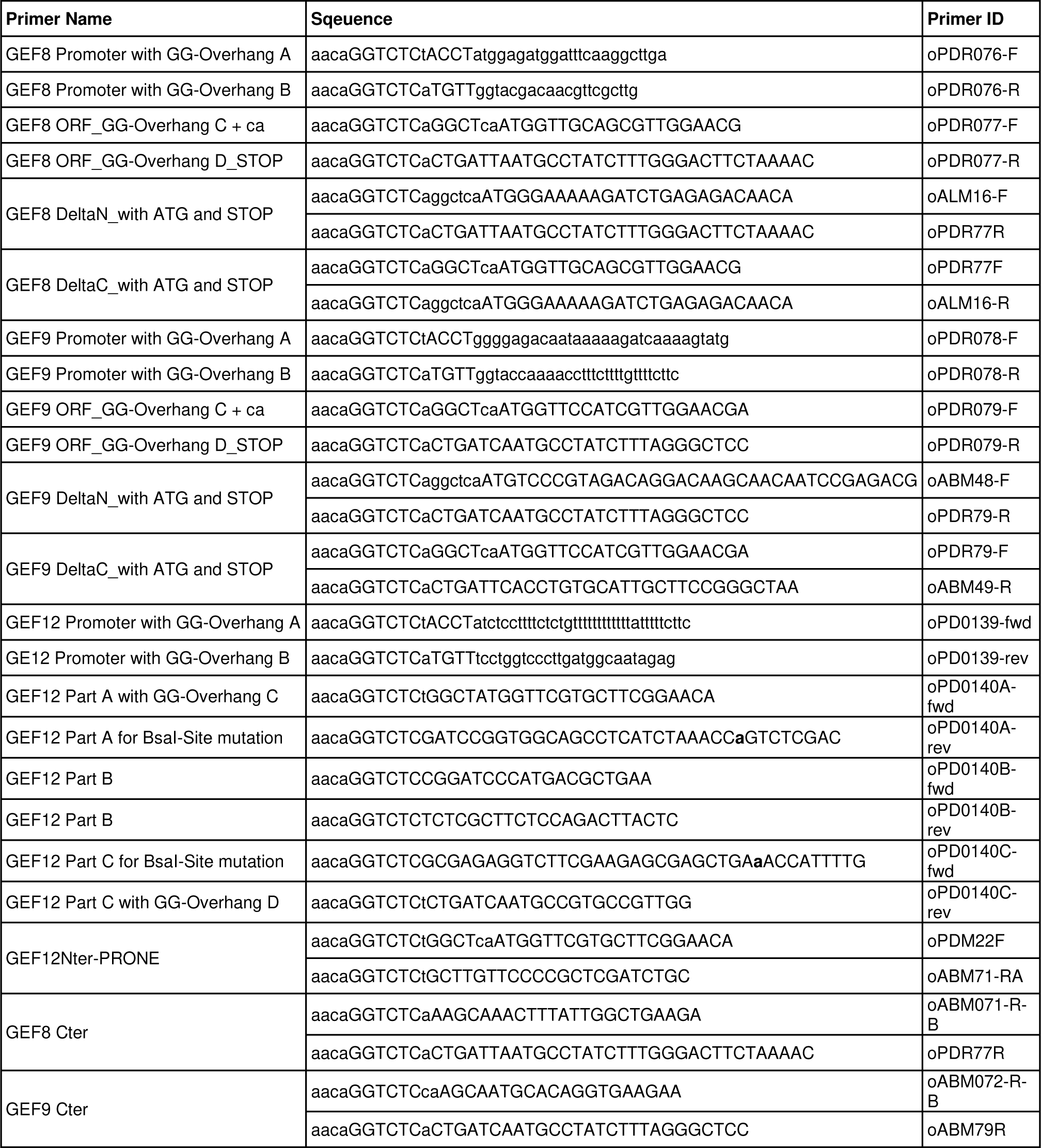
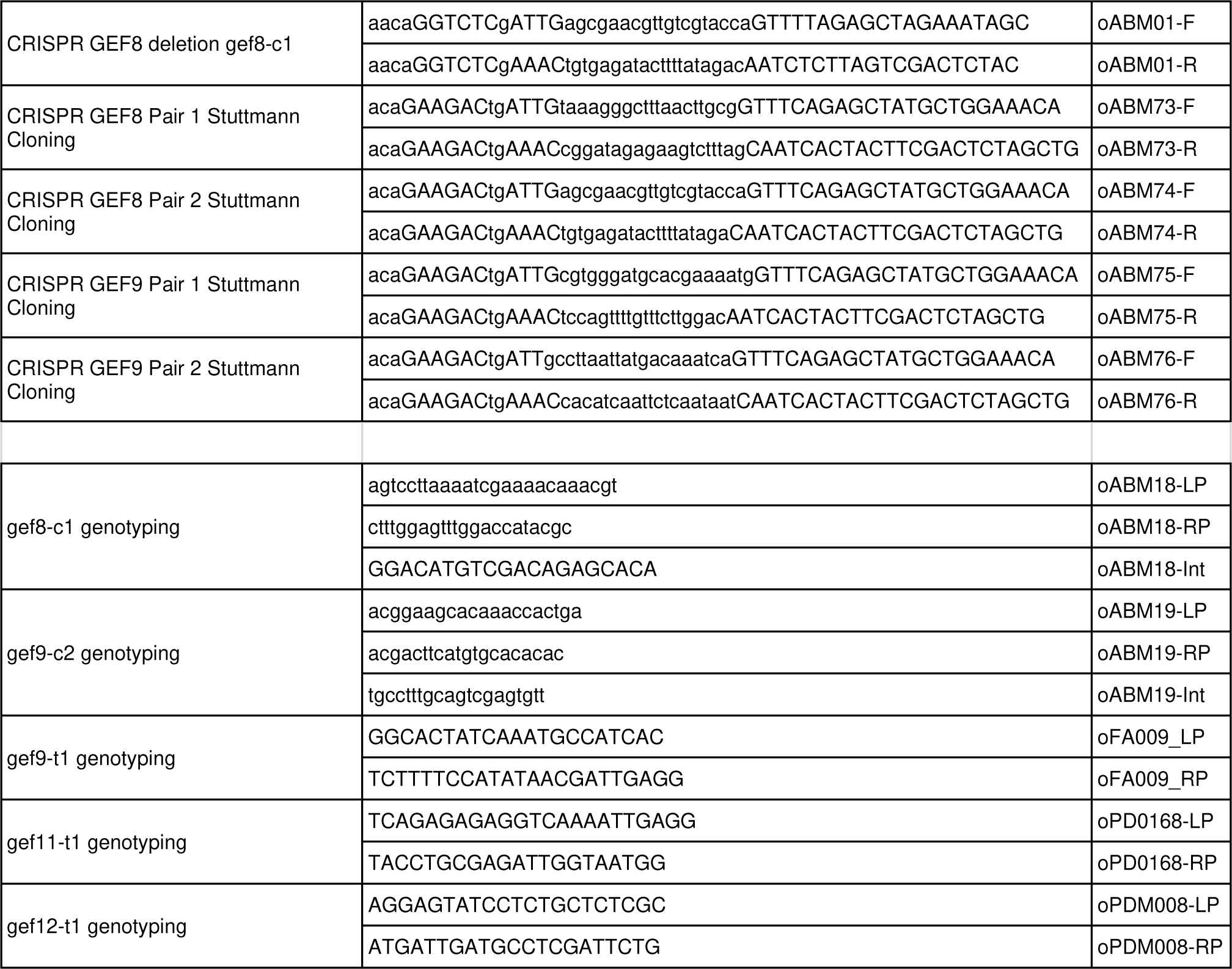
List of primers.

**Supplemental Table S2:** List of cloned expression vectors.

|  |  | Promoter | N-terminal Tag | ORF / CDS | C-terminal Tag | Terminator | Plant selection | Expression Vector | Bacterial selection marker | Plasmid ID |
| --- | --- | --- | --- | --- | --- | --- | --- | --- | --- | --- |
|  |  | Module A | Module B | Module C | Module D | Module E | Module F | Module Z |  |  |
| Green Gate Vector Cloning | **mCit-GEF8** | GEF8-Promoter | mCitrine w/ Linker | GEF8-ORF | Decoy | HSP18.2 | Basta | pGGZ003 | Spec/Strep | pPDR124 |
|  | **mCit-GEF9** | GEF9-Promoter | mCitrine w/ Linker | GEF9-ORF | Decoy | HSP18.2 | Basta | pGGZ003 | Spec/Strep | pPDR125 |
|  | **mCit-GEF12** | GEF12-Promoter | mCitrine w/ Linker | GEF12-ORF | Decoy | HSP18.2 | Basta | pGGZ003 | Spec/Strep | pPD0304 |
|  | **mSct-GEF8** | GEF8-Promoter | mScarlet w/ Linker | GEF8-ORF | Decoy | HSP18.2 | Hygromycine | pGGX000 | Kanamycince | pABM04 |
|  | **mSct-GEF9** | GEF9-Promoter | mScarlet w/ Linker | GEF9-ORF | Decoy | HSP18.2 | Hygromycine | pGGX000 | Kanamycince | pABM05 |
|  | **mSct-GEF12** | GEF12-Promoter | mScarlet w/ Linker | GEF12-ORF | Decoy | HSP18.2 | Hygromycine | pGGX000 | Kanamycince | pABM06 |
| | **mCit-GEF8ΔN** | GEF8-Promoter | mCitrine w/ Linker | GEF8-ORFΔN | Decoy | HSP18.2 | Basta | pGGYR0 | Spec/Strep | pABM93 |
| | **mCit-GEF8ΔC** | GEF8-Promoter | mCitrine w/ Linker | GEF8-ORFΔC | Decoy | HSP18.2 | Basta | pGGYR0 | Spec/Strep | pABM94 |
| | **mCit-GEF9ΔN** | GEF9-Promoter | mCitrine w/ Linker | GEF9-ORFΔN | Decoy | HSP18.2 | Basta | pGGYR0 | Spec/Strep | pABM76 |
| | **mCit-GEF9ΔC** | GEF9-Promoter | mCitrine w/ Linker | GEF9-ORFΔC | Decoy | HSP18.2 | Basta | pGGYR0 | Spec/Strep | pABM77 |
|  | **mCit-GEF12GEF8Cter** | GEF12-Promoter | mCitrine w/ Linker | GEF12-ORF+<br>GEF8Cter | Decoy | HSP18.2 | Basta | pGGYR0 | Spec/Strep | pABM83 |
|  | **mCit-GEF12GEF9Cter** | GEF12-Promoter | mCitrine w/ Linker | GEF12-ORF+<br>GEF9Cter | Decoy | HSP18.2 | Basta | pGGYR0 | Spec/Strep | pABM84 |
|  | **Gef12p::RIC4-CRIB:mCit** | GEF12-Promoter | Omega element for<br>enhanced translation | RIC4-CRIB | mCitrine w/<br>Linker | HSP18.2 | Basta | pGGYR0 | Spec/Strep | pPDM117 |
|  | **Lat52::RGeco1** | Lat52 promoter | Decoy | RGeco1 | Decoy | HSP18.4 | Hygromycine | pGGX000 | Kanamycince | pFA18 |
|  | **GEF8 GEF9 CRISPR** | GEF8 gRNAs-<br>pair 1 | GEF8 gRNAs- pair<br>2 | GEF9 gRNAs-<br>pair 1 | GEF9 gRNAs-<br>pair 2 | - | - | pDGE347 | Spec/Strep | pABM90 |

